# Knowledge, Attitudes and Practices of Hepatitis B Prevention and Immunization of Pregnant Women and Mothers in Viet Nam

**DOI:** 10.1101/470849

**Authors:** Thi T. Hang Pham, Thuy X. Nguyen, Dong T. Nguyen, Chau M. Luu, Bac D. Truong, Phu D. Tran, Samuel So

## Abstract

**Background and Aim:** Vietnam’s high burden of liver cancer is largely attributable to the high prevalence of chronic hepatitis B virus infection (HBV). Infection at birth due to mother-to-child (MTC) transmission is the most common cause of chronic HBV in Vietnam and increases the risk of liver cancer later in life. This study was undertaken to examine the knowledge, attitudes, and practices of pregnant women and mothers in Vietnam concerning HBV prevention and immunization.

**Methods:** A cross-sectional study was conducted in Quang Ninh and Hoa Binh provinces in 2017. A pre-designed questionnaire was administered to women when they received care at primary and tertiary maternal health clinics. Correct responses were summarized as knowledge scores. Data was analyzed using a multivariable regression model across participant demographics.

**Results:** Among the 404 women surveyed, 57.6% were pregnant and 42.4% were postpartum. Despite 73.5% of participants reporting having received information about HBV during their pregnancy, gaps in knowledge and misconceptions are evident. Overall, only 10.6% provided correct answers to all questions regarding HBV transmission routes and prevention measures. Around half of the participants incorrectly believed that HBV is transmitted through sneezing, contaminated water or sharing foods with chronic HBV patients. Although 96.4% of participants believed that HBV vaccination is necessary for infants, only 69.1% were willing to have their own child vaccinated within 24 hours. More than a third of participants expressed concern about having casual contacts or sharing foods with chronic HBV patients. In multivariable analysis, having received information about HBV during their pregnancy were consistently associated with better knowledge score for transmission, prevention and immunization. However, knowledge of women who received information about HBV during their pregnancy was still suboptimal.

**Conclusions:** The results highlight the need to prioritize educating pregnant women and mothers in future public health campaigns in order to increase knowledge, reduce misperception, and improve HBV vaccine coverage in Vietnam.

## INTRODUCTION

Liver cancer is the fourth most common cause of death from cancer and carried the second highest rate of absolute years of life lost among amongst cancers globally in 2016[1]. The 2016 Global Burden of Disease also estimates that HBV infection alone accounts for about 42% of liver cancer death.

Vietnam has the 6th highest incidence of liver cancer and the third highest rate of death from liver cancer in the world in 2018[2]. In 2018, liver cancer was the third leading cause of cancer death in Vietnam with a male-to-female ratio of 3.5 for age-standardized mortality. HBV accounts for close to half (46%) of the liver cancer deaths in Vietnam[3]. The reported prevalence of the HBV surface antigen (HBsAg) in the general population ranged from 15-20% [4-7]. Current estimates suggests 10.8% of the population or 9.6 million people in Vietnam are HBsAg positive and are living with chronic hepatitis B[5].

The spread and development of serious HBV sequelae can be effectively prevented through immunization with the hepatitis B vaccine. Infant hepatitis B vaccination was introduced to Vietnam in 1997 and expanded nationwide in 2002. To combat the significant risk of infection at birth due to MTC transmission, a birth dose administered within 24 hours after birth was added to the immunization schedule in 2003 [8]. The results of a nationwide survey comparing HBsAg prevalence in children born 2000-2003 to children born 2007-2008 showed a 2% reduction in prevalence. Additionally, infants vaccinated > 7 days after birth showed a 1.68% increase in HBsAg prevalence compared to those vaccinated within the 24 hour following the immunization guideline[9]. Such findings demonstrate the profound impact of both implementation and timely vaccine administration in reducing the risk of chronic hepatitis B infection.

Despite these encouraging results, birth-dose vaccination coverage has struggled to stay consistent since its implementation. Birth dose coverage dropped to its lowest rate in 2010 at 21.4%, rose to its highest in 2012 at 75% before dropping to 56% in 2013. The reporting of adverse events following immunization (AEFIs) in 2007 and 2013 that were blamed on newborn hepatitis B vaccination in conjunction the emergence of anti-vaccination movements may contribute to this volatility. The impact of a 19% drop in coverage was estimated to increase burden by 130,675 new chronic HBV infections and 25,197 HBV-associated deaths for children born in 2013 [10]. As perinatal transmission continues to be the major route of transmission in Vietnam, it is critical to initiate strategies to improve and sustain vaccine coverage rates. A pertinent strategy will be to recruit pregnant women and postnatal mothers to become active participants in their own health as well as advocates for the health of their children. There is currently a dearth in data regarding knowledge, attitudes, and sources of misconception regarding HBV prevention and care in Vietnamese mothers and pregnant women. The findings from this survey will be used to identify putative areas where targeted public health initiatives will be most effective in eliminating HBV and liver cancer in Vietnam.

## METHODS

### Study population

This was a cross sectional study recruiting a sample size of 404 pregnant or postnatal women at 16 selected maternal clinics at primary and tertiary hospitals in Hoa Binh and Quang Ninh provinces. Women who visited maternal health units were approached after their visit and invited to participate if they were pregnant or within 60 days postpartum. Trained data collectors administered a pre-designed questionnaire at maternal care clinics. Written consent form was obtained from the participants before interviewing.

### Questionnaire

The questionnaire was developed in Vietnamese by the Asian Liver Center at Stanford University based on its past experience with administering HBV knowledge surveys in other populations. The questionnaire consisted of four sections: i) demographic and personal HBV-related health history; ii) disease burden and consequences; iii) transmission routes and prevention measures; iv) postnatal mother HBV vaccination practices. The first section surveyed participant demographics and personal HBV-related history. The second section on disease burden, consequences and risk factors tested for knowledge regarding national disease prevalence and consequences of chronic HBV infection. The third section on transmission routes and prevention measures examined knowledge regarding modes of transmission, prevention measures, infant vaccination and attitudes towards patients with chronic HBV. The fourth section was completed exclusively by postnatal women and elucidated what HBV vaccination protocol was followed in their most recent birth.

### Statistical methods

Descriptive statistics were generated from variables in the data obtained from the 404 pregnant and postnatal women who completed the survey. A correct response to each question received one point and incorrect or missing responses received no points. The knowledge score was calculated based on the sum of correct answers to the 12 transmission and prevention questions and 4 immunization questions. Association between demographic factors, access to HBV education during pregnancy and knowledge scores was estimated using a multivariable regression model.

## RESULTS

Demographics and pregnancy characteristics of the study population (N = 404) are presented in Table 1. Of the women surveyed, 57.6% were pregnant and 42.4% in postpartum period. Participants’ age range from 17 to 45 years (median age: 27 years, mean age: 27 years) with the majority (86%) falling within the 20-34 years age group. 74.4% have completed high school level education or higher. The most common occupations were farming (23.3%), housewife (22.2%), and clerk or administrative positions (20.8%). Among the women surveyed, 14.2% had household per capita income that falls below the 2015 national poverty line (400,000 Vietnam Dong or 17 USD each month). Location of antenatal care was primarily at province level hospitals (46.8%) with the remaining approximately evenly split between district (27.1%) and commune health facilities (22.0%). Only 2.3% receive antenatal care at private clinics.

**Table 1.**
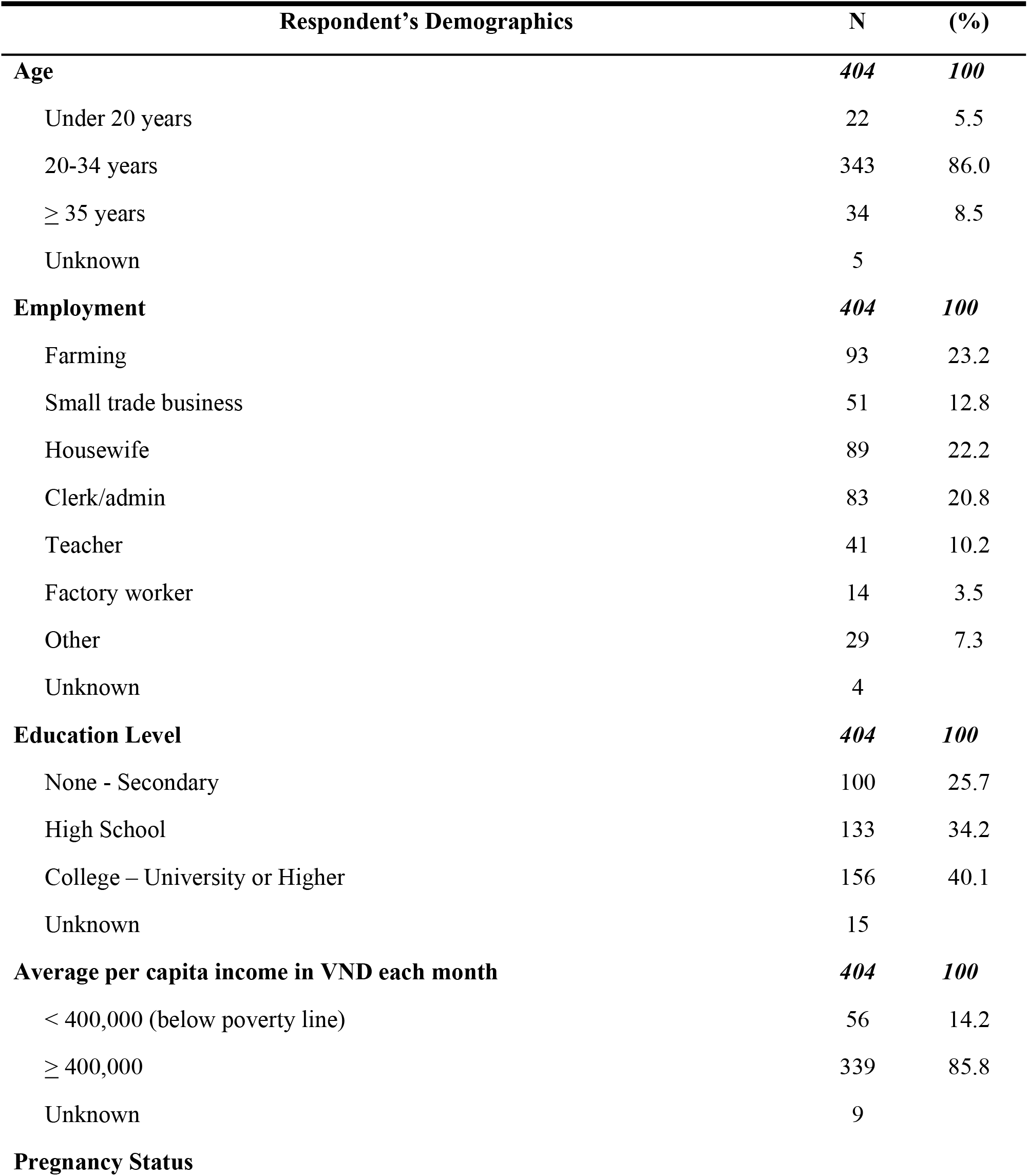
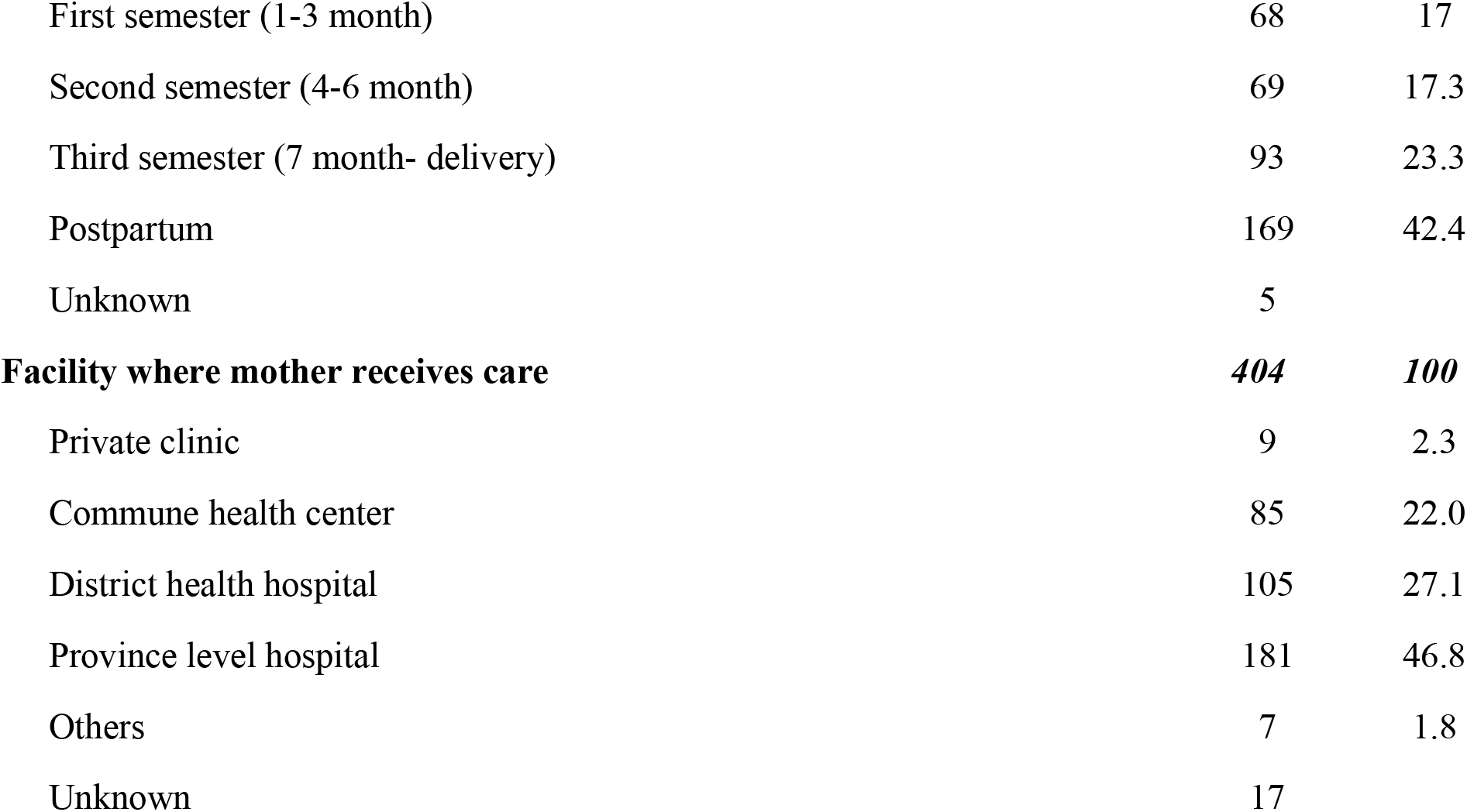
Demographic factors of survey participants (N = 404 pregnant/postnatal women)

| **Respondent's Demographics** | **N** | **(%)** |
| --- | --- | --- |
| **Age** | **404** | **100** |
| Under 20 years | 22 | 5.5 |
| 20-34 years | 343 | 86.0 |
| ≥ 35 years | 34 | 8.5 |
| Unknown | 5 |  |
| **Employment** | **404** | **100** |
| Farming | 93 | 23.2 |
| Small trade business | 51 | 12.8 |
| Housewife | 89 | 22.2 |
| Clerk/admin | 83 | 20.8 |
| Teacher | 41 | 10.2 |
| Factory worker | 14 | 3.5 |
| Other | 29 | 7.3 |
| Unknown | 4 |  |
| **Education Level** | **404** | **100** |
| None - Secondary | 100 | 25.7 |
| High School | 133 | 34.2 |
| College – University or Higher | 156 | 40.1 |
| Unknown | 15 |  |
| **Average per capita income in VND each month** | **404** | **100** |
| < 400,000 (below poverty line) | 56 | 14.2 |
| ≥ 400,000 | 339 | 85.8 |
| Unknown | 9 |  |
| **Pregnancy Status** |  |  |
| First semester (1-3 month) | 68 | 17 |
| Second semester (4-6 month) | 69 | 17.3 |
| Third semester (7 month- delivery) | 93 | 23.3 |
| Postpartum | 169 | 42.4 |
| Unknown | 5 |  |
| **Facility where mother receives care** | **404** | **100** |
| Private clinic | 9 | 2.3 |
| Commune health center | 85 | 22.0 |
| District health hospital | 105 | 27.1 |
| Province level hospital | 181 | 46.8 |
| Others | 7 | 1.8 |
| Unknown | 17 |  |

### General knowledge and access to information about HBV

Knowledge about HBV prevalence and its serious consequences was inadequate amongst pregnant women and mothers. About two thirds were not aware of the high prevalence of chronic hepatitis B infection in Vietnam. Only 58.8% of participants were aware that chronic HBV can cause serious consequences such as liver cirrhosis, liver failure, liver cancer, or premature death (Table 2). 69.5% of participants reported that they received information about HBV during their pregnancy. 81.7% reported that they have previously received information about the benefit of HBV vaccine for infants. Healthcare workers were the primary source of this information (90.1%) followed by equal contribution from common public outreach methods such as flyers, newspapers, radio, television, and the internet (Table 2).

**Table 2:**
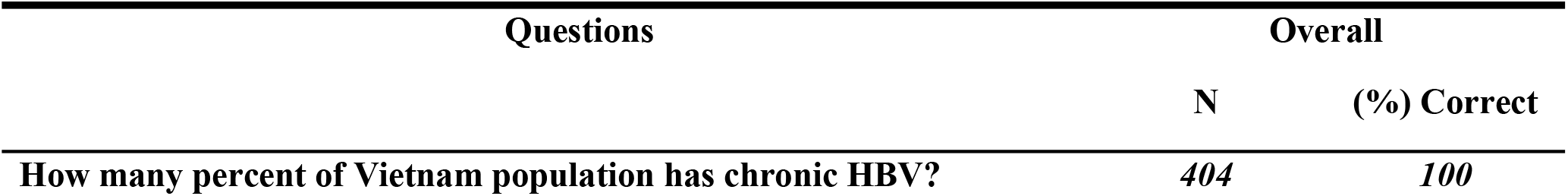
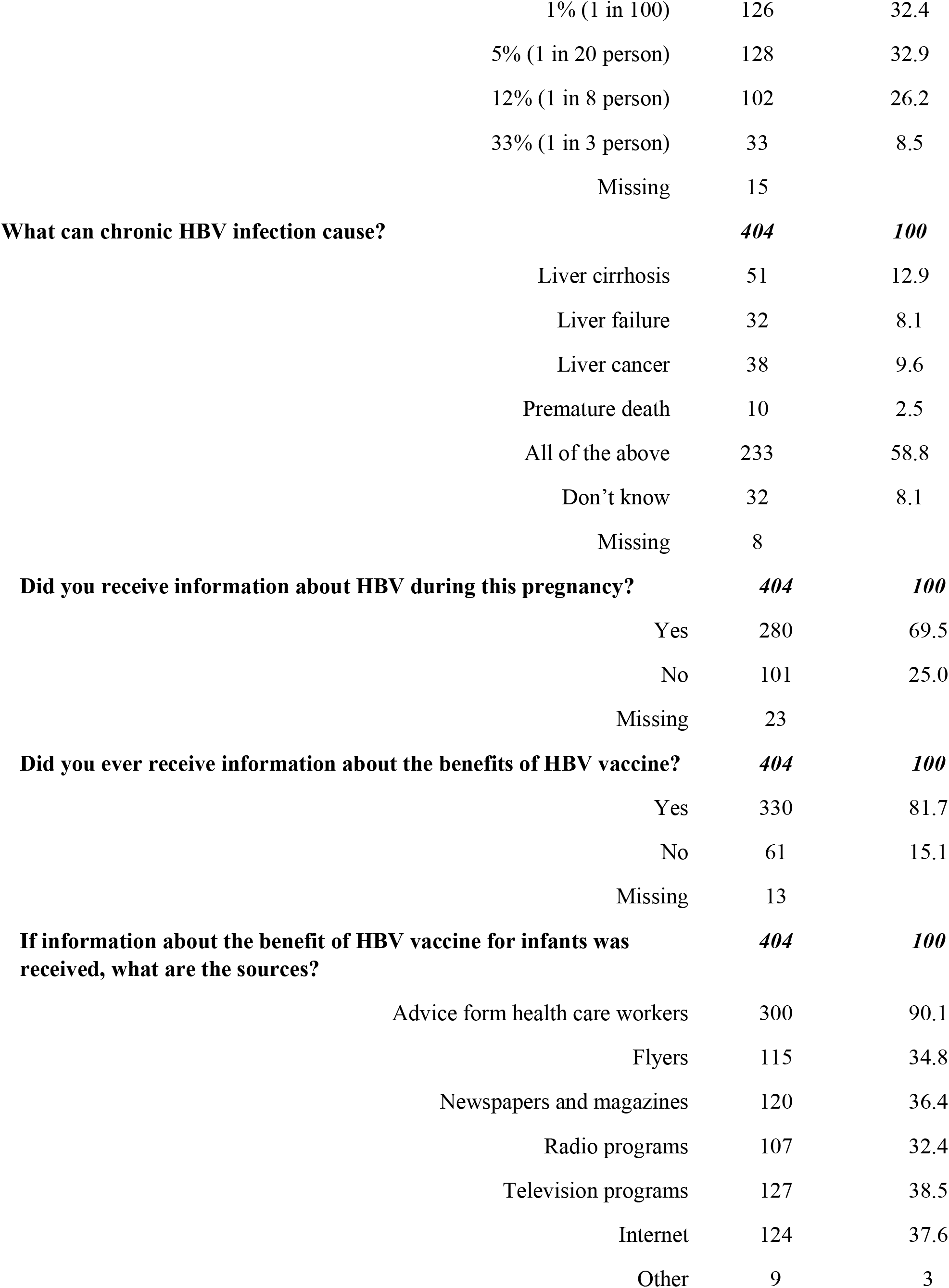
General knowledge and access to information about HBV (N = 404 pregnant/postnatal women)

| Questions | Overall |  |
| --- | --- | --- |
|  | N | (%) Correct |
| How many percent of Vietnam population has chronic HBV? | 404 | 100 |

|  |  |  |  |
| --- | --- | --- | --- |
|  | 1% (1 in 100) | 126 | 32.4 |
|  | 5% (1 in 20 person) | 128 | 32.9 |
|  | 12% (1 in 8 person) | 102 | 26.2 |
|  | 33% (1 in 3 person) | 33 | 8.5 |
|  | Missing | 15 |  |
| **What can chronic HBV infection cause?** |  | **404** | **100** |
|  | Liver cirrhosis | 51 | 12.9 |
|  | Liver failure | 32 | 8.1 |
|  | Liver cancer | 38 | 9.6 |
|  | Premature death | 10 | 2.5 |
|  | All of the above | 233 | 58.8 |
|  | Don't know | 32 | 8.1 |
|  | Missing | 8 |  |
| **Did you receive information about HBV during this pregnancy?** |  | **404** | **100** |
|  | Yes | 280 | 69.5 |
|  | No | 101 | 25.0 |
|  | Missing | 23 |  |
| **Did you ever receive information about the benefits of HBV vaccine?** |  | **404** | **100** |
|  | Yes | 330 | 81.7 |
|  | No | 61 | 15.1 |
|  | Missing | 13 |  |
| **If information about the benefit of HBV vaccine for infants was received, what are the sources?** |  | **404** | **100** |
|  | Advice from health care workers | 300 | 90.1 |
|  | Flyers | 115 | 34.8 |
|  | Newspapers and magazines | 120 | 36.4 |
|  | Radio programs | 107 | 32.4 |
|  | Television programs | 127 | 38.5 |
|  | Internet | 124 | 37.6 |
|  | Other | 9 | 3 |

### Knowledge and attitude regarding HBV transmission and prevention

Out of 12 questions about HBV transmission and prevention knowledge, the mean score was 8.2 ± 2.67 (mean +-SD) and the median score was 8.0. Study participants were largely aware that HBV can be transmitted through mother-to-child (85.1%), unprotected sex (75.3%), and blood transfusions (86.7%). However, there were common misconceptions that HBV can be transmitted through sneezing or coughing (58.5%), contaminated water (55.2%), and eating with or sharing food with chronic HBV patients (47.1%) (Table 3). Only 43 out of 404 participants (10.6%) provided correct answers to all 12 questions regarding HBV transmission routes and prevention measures.

Participants’ knowledge about preventive measures of HBV infection was consistent with their level of knowledge about transmission routes. The percentage of participants who were aware of that HBV can be prevented by receiving the hepatitis B vaccine, not reusing or share needle/syringes and using condom were 93.5%, 91.5% and 79.3% respectively. Because of the misconceptions that HBV can be transmitted through contaminated water or sharing food, only 28.6% participants provided correct answer to whether thorough cooking and cleaning of food can prevent HBV transmission or not. 40.6% provided correct answer to whether avoiding sharing food and utensils with chronic HBV individuals can prevent HBV transmission (Table 3).

**Table 3.**
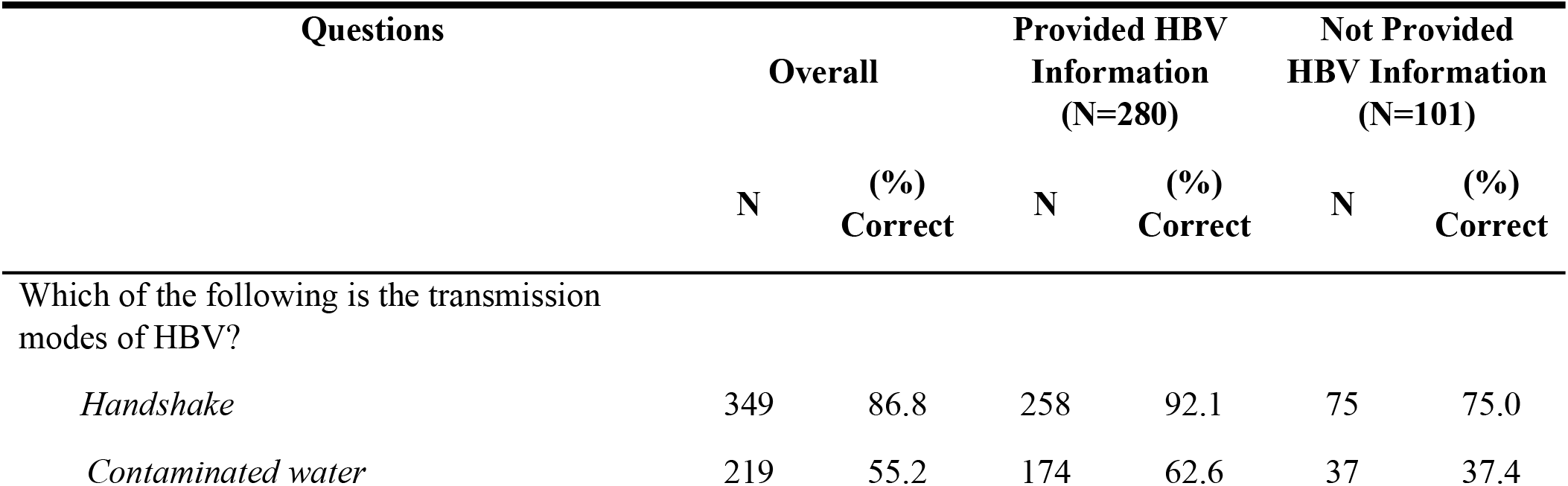
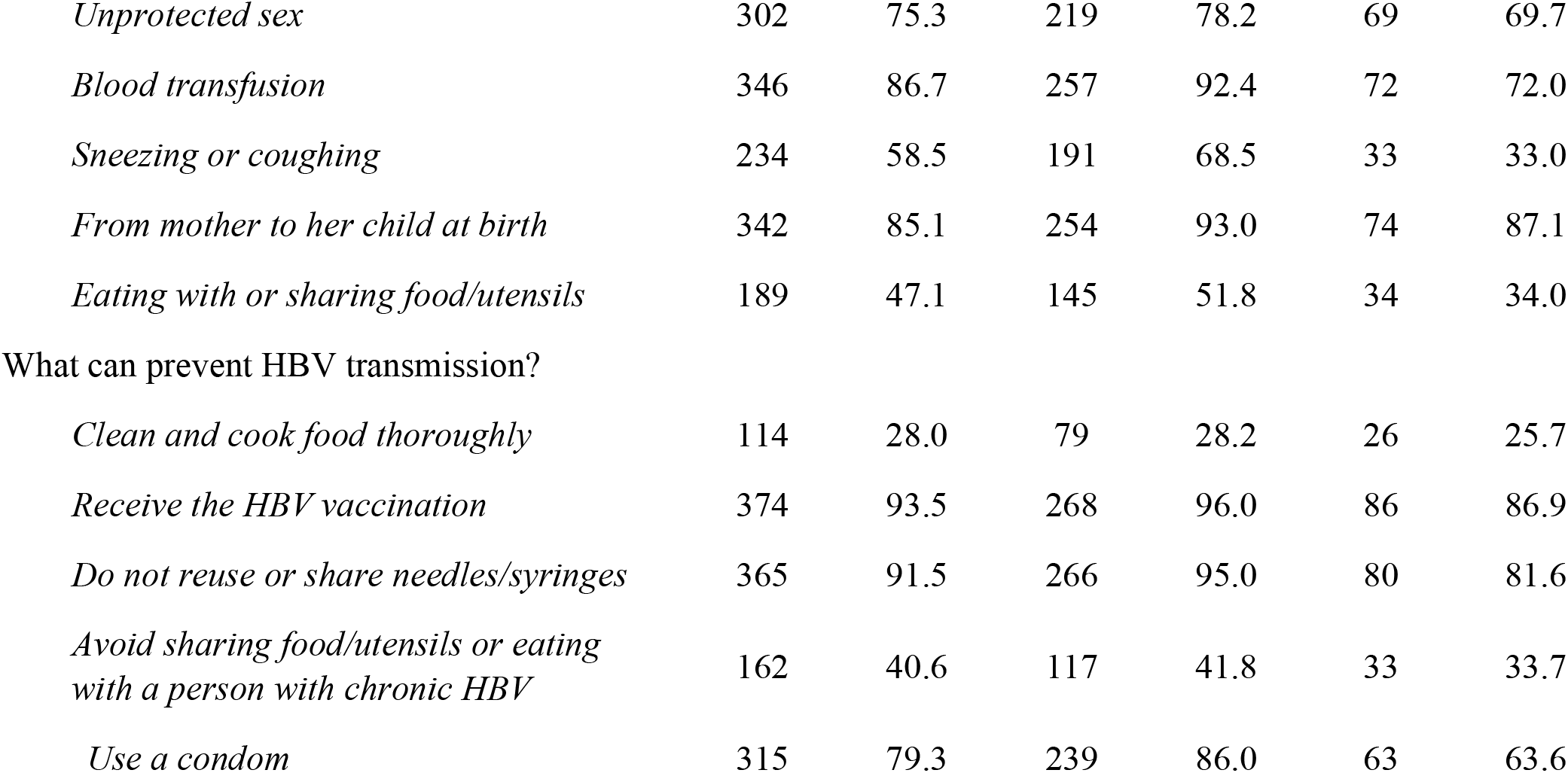
Distribution of correct responses to HBV transmission and prevention related questions based on receipt of information during pregnancy (N = 404 pregnant/postnatal women)

| Questions | Overall |  | Provided HBV Information (N=280) |  | Not Provided HBV Information (N=101) |  |
| --- | --- | --- | --- | --- | --- | --- |
|  | N | (%) Correct | N | (%) Correct | N | (%) Correct |
| Which of the following is the transmission modes of HBV? |  |  |  |  |  |  |
| *Handshake* | 349 | 86.8 | 258 | 92.1 | 75 | 75.0 |
| *Contaminated water* | 219 | 55.2 | 174 | 62.6 | 37 | 37.4 |
| *Unprotected sex* | 302 | 75.3 | 219 | 78.2 | 69 | 69.7 |
| *Blood transfusion* | 346 | 86.7 | 257 | 92.4 | 72 | 72.0 |
| *Sneezing or coughing* | 234 | 58.5 | 191 | 68.5 | 33 | 33.0 |
| *From mother to her child at birth* | 342 | 85.1 | 254 | 93.0 | 74 | 87.1 |
| *Eating with or sharing food/utensils* | 189 | 47.1 | 145 | 51.8 | 34 | 34.0 |
| What can prevent HBV transmission? |  |  |  |  |  |  |
| *Clean and cook food thoroughly* | 114 | 28.0 | 79 | 28.2 | 26 | 25.7 |
| *Receive the HBV vaccination* | 374 | 93.5 | 268 | 96.0 | 86 | 86.9 |
| *Do not reuse or share needles/syringes* | 365 | 91.5 | 266 | 95.0 | 80 | 81.6 |
| *Avoid sharing food/utensils or eating with a person with chronic HBV* | 162 | 40.6 | 117 | 41.8 | 33 | 33.7 |
| *Use a condom* | 315 | 79.3 | 239 | 86.0 | 63 | 63.6 |

About a third of the surveyed pregnant women and mothers had concerns about having casual contact (32.9%), working with or sharing food with chronic HBV patients (38.1%). Moreover, 42.5% of respondents expressed having concerns if their child was in the same class with a child with chronic HBV infection (Table 4)

**Table 4.** Attitudes towards HBV transmission (N = 404 pregnant/postnatal women)

| Question | Overall<br>N=404 |  | Provided<br>HBV<br>Information<br>(N=280) |  | Not Provided<br>HBV<br>Information<br>(N=101) |  |
| --- | --- | --- | --- | --- | --- | --- |
|  | N | (%) | N | (%) | N | (%) |
| Do you have concerns about interacting with chronic HBV patients in the following ways? (N = yes) |  |  |  |  |  |  |
| Casual contact or working together in the same office | 130 | 32.9 | 88 | 31.4 | 40 | 39.6 |
| Eating with (sharing food or utensils) | 151 | 38.1 | 101 | 36.1 | 44 | 42.3 |
| If your child is in the same class with a chronic HBV kid | 168 | 42.5 | 105 | 37.5 | 51 | 50.5 |

In multivariable analysis, having received information about HBV during pregnancy was the only factor independently associated with transmission and prevention score. Age, number of children, family per-capital income and education level were not associated with transmission and prevention knowledge score (Table 5). While women who received information about HBV during pregnancy provided higher percentages of correct answers to almost all individual questions related to HBV transmission and prevention compared to those who did not, their knowledge were still suboptimal. Only 62.6%, 68.5% and 51.8% respectively provided correct answers to whether HBV can be transmitted through contaminated water, coughing/sneezing and sharing foods. Only 28.2% provided correct answer to whether cleaning and cooking food thoroughly can prevent HBV transmission or not; 41.8% provided correct answers to whether avoiding sharing food/utensils or eating with a person with chronic HBV or not (Table 3).

**Table 5.**
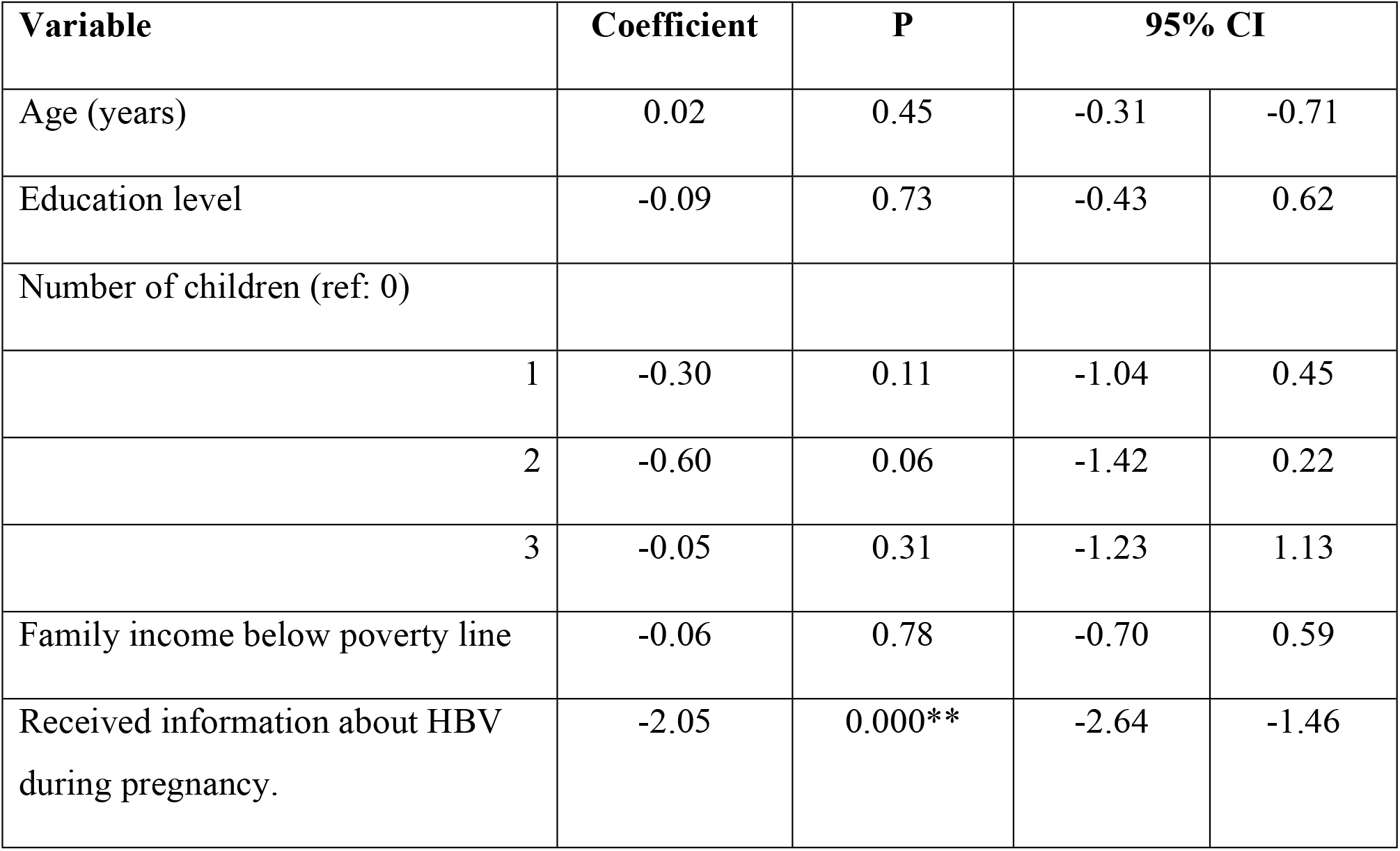
Analysis of factors associated with transmission and prevention knowledge scores. Variable Coefficient P 95% CI.

### HBV screening and immunization knowledge and attitude

Out of 4 questions about HBV screening and immunization, the mean knowledge score for was 3.0 ± 0.98 (mean ± SD) and the median was 3.0. Majority of surveyed women believed that pregnant women need to be tested for HBV (83.2%), vaccination is necessary for infants (96.4%) and the best time to provide a healthy and stable child the first dose of HBV vaccine is within 24 hours after birth (80.9%). However, only 54.5% of participants knew that infants born to chronic HBV mothers should receive the first dose of the hepatitis B vaccine and the HBIG shot within 12 hours of birth followed by completion of the vaccine series to prevent mother to child infection (Table 6).

**Table 6:** Distribution of correct responses to HBV screening and immunization knowledge questions based on receipt of information (N = 404 pregnant/postnatal women)

| Questions | Overall |  | Provided HBV Information (N=280) |  | Not Provided HBV Information (N=101) |  |
| --- | --- | --- | --- | --- | --- | --- |
|  | N | (%) Correct | N | (%) Correct | N | (%) Correct |
| As a pregnant woman, do you need to be tested for HBV? | 331 | 83.2 | 262 | 93.9 | 60 | 61.2 |
| Is the HBV vaccination is necessary for your infant? | 346 | 96.4 | 251 | 98.0 | 81 | 96.4 |
| When is best time to provide a healthy and stable child the first dose of HBV vaccine? | 322 | 80.9 | 233 | 83.5 | 74 | 75.5 |
| If a pregnant woman has chronic HBV, what measure could protect the newborn from becoming infected? | 207 | 54.5 | 148 | 55.0 | 49 | 53.2 |

While surveyed women had fairly good knowledge about hepatitis B vaccination, their confidence in giving their own children the hepatitis B vaccine birth dose was lower. Only 69.1% believed that the vaccine is very safe and would be willing to vaccinate their own healthy baby within the first 24 hours of birth. 66.4% of mothers said they would definitely be willing to have their own child vaccinated within 24 hours after birth even after reassurance of the hepatitis B vaccine’s safety by a physician

**Table 7.** Attitudes towards HBV vaccine birth dose in pregnant and postnatal women (N = 404 pregnant/postnatal women)

| Question | N | (%) |
| --- | --- | --- |
| If your newborn is healthy and stable, do you think it is safe to give your baby HBV vaccine in the first 24 hours after birth? |  |  |
| Very safe | 279 | 69.1 |
| Maybe safe | 77 | 19.1 |
| Not very safe | 26 | 6.4 |
| Missing | 22 |  |
| If your doctor tells you that HBV vaccine is safe to given to newborn, would you be willing to get your child HBV vaccine? |  |  |
| Definitely | 263 | 66.4 |
| Maybe | 106 | 26.8 |
| Not sure | 23 | 5.8 |
| No | 4 | 1.0 |
| Unknown | 8 |  |

In multivariate analysis, characteristics independently associated with higher screening and immunization knowledge scores included ages of participants and having received information about HBV during pregnancy (Table 8). Younger women tend to have lower screening and immunization score compared to their peers of older age. Number of children, family per-capital income and mother education were not associated with screening and immunization knowledge score.

**Table 8.** Analysis of factors associated with HBV screening and immunization knowledge scores.

| Variable | Coefficient | P | 95% CI |  |
| --- | --- | --- | --- | --- |
| Age(years) | 0.03 | 0.000* | 0.01 | 0.05 |
| Education | 0.05 | 0.73 | -0.13 | 0.24 |
| Number of children (ref: 0) |  |  |  |  |
| 1 | -0.01 | 0.94 | -0.25 | 0.27 |
| 2 | -0.14 | 0.06 | -1.42 | 0.22 |
| 3 | -0.19 | 0.37 | -0.61 | 0.23 |
| Family per-capita income | -0.11 | 0.37 | -0.34 | 0.12 |
| Received information about HBV during pregnancy. | -0.57 | 0.000** | -0.78 | -0.37 |

### HBV screening and immunization practice

In the subgroup of 203 postnatal women surveyed, 68.4% reported they received hepatitis B testing during their current or most recent pregnancy. Among them, 20% reported having positive results and 16% were unsure of their results. 71.6% reported the newborn were administered the first dose of the hepatitis B vaccine within 24 hours of birth. 13.7% were vaccinated between 24 to 48 hours after birth and 14.7% did not receive any vaccine until 1 month of age. When asked why the infants were not vaccinated within 24 hours of birth, the following responses were given: mother did not think it was safe (30.9%); no vaccine available (17.7%); child was sick (14.7%); mother did not think it was necessary (13.2); doctor said it was not necessary (10.3%) and child has low birth weight (2.9%) (Table 8).

**Table 8.**
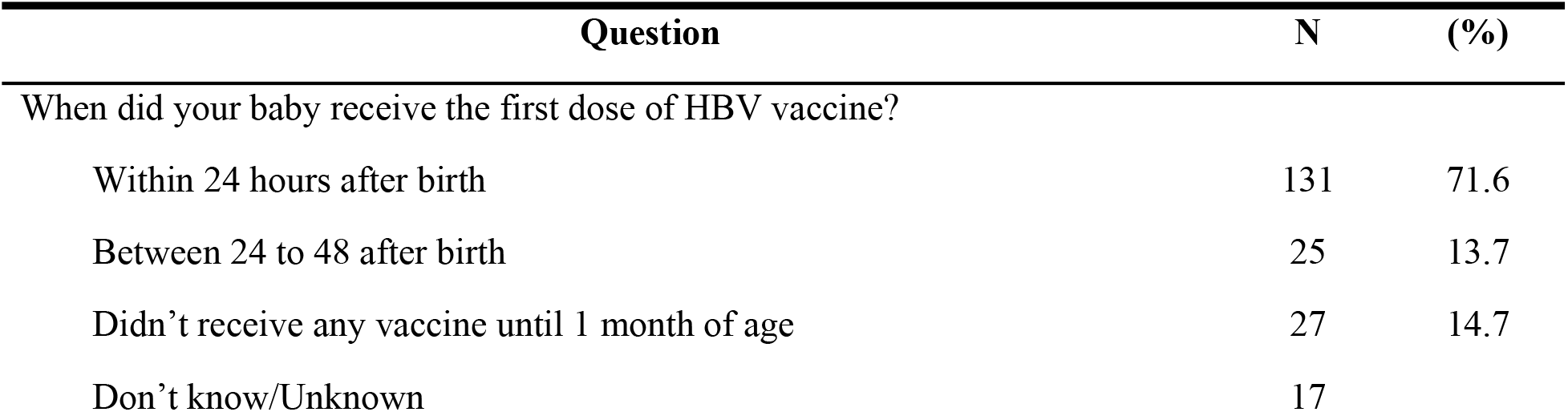
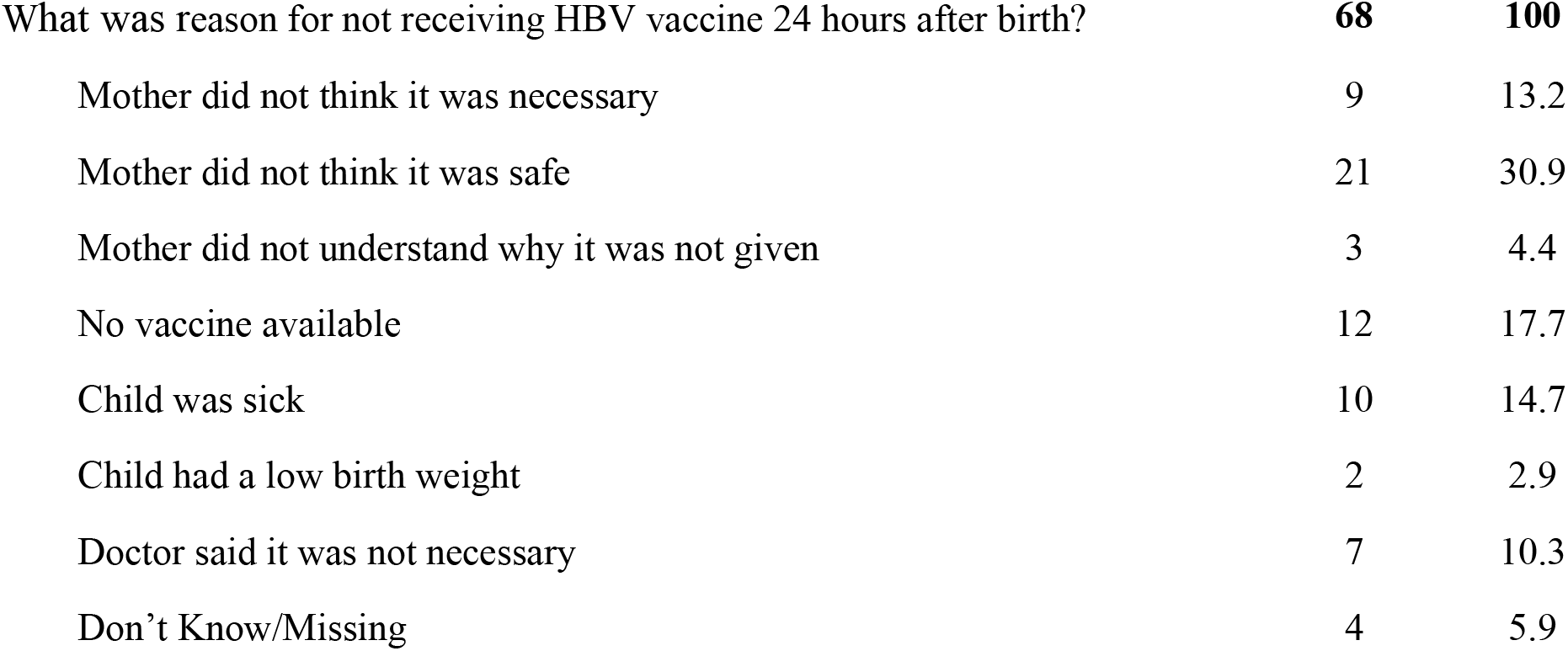
Local HBV birth dose vaccination practices (N = 203 postnatal women)

In multivariable analysis, having received information about benefits of HBV vaccination for newborn before and delivering at provincial health clinics were independently associated with whether the newborns received the hepatitis B birth dose or not. Age, education level, number of children, family per-capital income and knowledge score on HBV screening and immunization were not associated with whether the newborns received the hepatitis B birth dose or not (Table 9)

**Table 9.** Analysis of factors associated with newborn receiving HBV vaccine birth dose.

| Variable | Coefficient | P | 95% CI |  |
| --- | --- | --- | --- | --- |
| Age(years) | -0.00 | 0.82 | -0.15 | 0.12 |
| High school or higher education | -0.13 | 0.86 | -0.16 | 0.13 |
| Number of children (ref: pregnant women) |  |  |  |  |
| 1 | -0.51 | 0.10 | -1.13 | 0.92 |
| 2 | -0.51 | 0.08 | -1.09 | 0.68 |
| 3 | -0.46 | 0.11 | -1.04 | 0.11 |
| Per-capita income under poverty line | -0.06 | 0.50 | -0.24 | 0.11 |
| Having received information about benefits of HBV vaccination for newborn | -0.33 | 0.000* | -0.52 | -0.15 |
| Place where delivery is taking place (ref: commune health center) |  |  |  |  |
| Provincial health clinics | -0.29 | 0.03*** | -0.55 | -0.22 |
| District health clinics | -0.19 | 0.12 | -0.43 | 0.05 |
| Private clinic | -0.45 | 0.80 | -0.39 | 0.30 |
| Others | -0.14 | 0.54 | -0.60 | 0.31 |
| HBV screening and immunization knowledge | 0.07 | 0.1 | -0.13 | 0.15 |

## DISCUSSION

This survey showed that pregnant women and mothers in Vietnam’s Quang Ninh and Hoa Binh provinces lacked knowledge regarding HBV transmission and prevention regardless of age, education, economic condition and childbearing status. Limited knowledge regarding HBV among pregnant women is consistent with findings from previous studies in other high epidemic countries [11-13]. Multivariable analysis showed that mothers who received HBV information during pregnancy consistently had better knowledge regarding HBV transmission, prevention and immunization than who did not. However, this knowledge among women who received HBV information during pregnancy was still sub-optimal. Our findings emphasize a need to implement education programs targeting women of childbearing ages with basic HBV information. It is also necessary to review existing antenatal educational programs and materials to ensure that key messages are effectively conveyed to the target audiences.

This study also revealed significant stigma associated with people having chronic HBV in Vietnam. About a third of participants expressed concerned about eating with, sharing food, casual contact or working in the same office with chronic HBV patients. A higher proportion (42.5%) expressed concerns if their children were in the same class with a child with chronic HBV infection. One explanation could be because approximately half of the women surveyed believed that HBV can be transmitted through sneezing or coughing, contaminated food and water, or eating or sharing food with chronic HBV patients. The stigma and pattern of knowledge deficits observed in this study regarding HBV was similar to a previous study among adult residents in Ho Chi Minh city in which 55% had the mistaken impression that HBV can be spread by sharing eating utensils and 61% felt that persons with chronic HBV infection put others at risk [14]. It is widely recognized that HBV related stigma can negatively affect health behaviors related to screening, preventive, diagnostic and treatment for HBV infection [15]. Further research into this area to understand the magnitude, underlying reasons and its impact is necessary to evaluate effective interventions to improve awareness and tackle stigma in HBV in Vietnam.

Knowledge of pregnant and mothers about hepatitis B vaccine was fairly good but their confidence in having their own baby vaccinated at birth was lower. Mother did not think hepatitis B vaccine was totally safe was the primary reasons given by mothers in this study for why their newborn did not receive HBV birth dose (32.8%). Residual effects from the 2007 and 2013 AEFIs may explain pregnant women’s hesitant attitude towards infant hepatitis B vaccination. It is important to note that this study was conducted back to back with a survey on HBV knowledge and practices of health care workers (HCWs) in the same clinics in which only 60.8% of surveyed HCWs felt confident that the hepatitis B vaccine is safe (unpublished data). This low confidence in HBV vaccine safety among HCWs may significantly contributed to high hesitance among pregnant women because they are the main source from which pregnant women received hepatitis B vaccine related information. These together underscored a critical need to address this concern to re-establish and sustain confidence in hepatitis B vaccine at a wider scope, targeting HCWs, pregnant women and general community.

### Study Limitations

This study was conducted in two Northern provinces of Vietnam with low HBV birth dose coverage at the study point; thus, the results may not be applicable to other parts of the country. In addition, the self-reported HBV screening and immunization practices by study participants could not be validated.

### Conclusion

This study showed that pregnant women and mothers have insufficient knowledge regarding HBV infection regardless of age, education, economic condition, childbearing status and historical exposure to HBV information during pregnancy. Misconceptions about HBV transmission through contaminated water, sharing foods and casual contacts were common and perpetuates the stigma associated with chronic HBV infection. Although most participants were aware of benefits of hepatitis B vaccine, concerns about vaccine safety for newborn was prevalent. These emphasized a need to enhance public health education efforts to improve hepatitis B knowledge among women in reproductive age and in the antenatal period, and to demystify issues surrounding HBV transmission and vaccine safety to improve hepatitis B birth dose vaccination rate and eliminate mother to child transmission.

